# The LIM-Only Protein FHL2 is involved in autophagy to regulate the development of skeletal muscle cell

**DOI:** 10.1101/459800

**Authors:** Zihao Liu, Shunshun Han, Yan Wang, Can Cui, Qing Zhu, Xiaosong Jiang, Chaowu Yang, Huarui Du, Chunlin Yu, Qingyun Li, Haorong He, Xiaoxu Shen, Yuqi Chen, Yao Zhang, Lin Ye, Zhichao Zhang, Diyan Li, Xiaoling Zhao, Huadong Yin

**Author notes:** These authors contributed equally to this work (Zihao Liu, Shunshun Han, Yan Wang, Can Cui). Corresponding author: **Huadong Yin**, Farm Animal Genetic Resources Exploration and Innovation Key Laboratory of Sichuan Province, Sichuan Agricultural University, Chengdu, Sichuan 611130, PR China.

## Abstract

Four and a half LIM domain protein 2 (FHL2) is a LIM domain protein expressed in muscle tissue whose deletion is causative of myopathies. Although FHL2 has a confirmed important role in muscle development, its autophagy-related function in muscle differentiation has not been fully determined. To explore the role of FHL2 in autophagy-related muscle regulation, FHL2-silenced and -overexpressing C2C12 mouse cells were examined. Immunofluorescence and co-immunoprecipitation assay findings showed that FHL2 silencing reduced LC3-Ⅱ protein expression and the amount of LC3 that co-immunoprecipitated with FHL2, indicating that FHL2 interacts with LC3-Ⅱ in the formation of autophagosomes. Moreover, the expression of muscle development marker genes such as MyoD1 and MyoG was lower in FHL2-silenced C2C12 cells but not in FHL2-overexpressing C2C12 cells. Electron microscopy analysis revealed large empty autophagosomes in FHL2-silenced myoblasts, while flow cytometry suggested that FHL2 silencing made cells more vulnerable to staurosporine-induced cell death. In conclusion, we propose that FHL2 interacts with LC3-Ⅱ in autophagosome formation to regulate the development of muscle cells.

## Introduction

Four and a half Lim domain protein 2 (FHL2) belongs to the FHL protein family which contains five members: FHL1, FHL2, FHL3, FHL4, and ACT [1]. Despite their high level of conservation among species, their expression levels differ from each other between tissues. FHL1, FHL2, and FHL3 are mainly expressed in skeletal and heart muscle [2], whereas FHL4 and ACT are highly expressed in the testis [3]. FHL2 has been shown to have a dual function in interacting with the cytoplasmic domain of several integrin chains [4], and also as a transcriptional coactivator of the androgen receptor [5]. Although FHL2 plays an important role in muscle development and its deletion was reported to lead to the development of myopathies [2, 6-10], the details of its function in skeletal muscle development are unclear.

Autophagy is the major intracellular degradation system by which cytoplasmic materials are delivered to and degraded in the lysosome, which also serves as a dynamic recycling system that produces new building blocks and energy for cellular renovation and homeostasis [11, 12]. Recently, the relationship between autophagy and muscle cell development has been shown to play an important role in muscle mass maintenance and integrity [13, 14]. As the main proteolytic system that controls protein degradation in skeletal muscle cells, the autophagy lysosome is activated in a number of catabolic disease states that lead to muscle loss and weakness [15, 16]. Excessive activation of autophagy aggravates muscle wasting by removing some cytoplasm, proteins, and organelles; conversely, the inhibition or alteration of autophagy can contribute to myofiber degeneration and weakness in muscle disorders [17]. Additionally, autophagy protects against apoptosis during myoblast differentiation [18].

Recently, a relationship between FHL2 and autophagy was identified, with involvement of the FHL2-activated nuclear factor-κB pathway reported in particulate matter 2.5-induced autophagy in mouse aortic endothelial cells [19]. Muscle Lim protein (MLP)/CSRP3 was reported to interact with microtubule-associated protein 1 light chain 3 (LC3) to regulate the differentiation of myoblasts and facilitate autophagy [20]. Moreover, Sabatelli et al found that the aggresome–autophagy pathway was involved in the pathophysiological mechanism underlying the muscle pathology of the C150R mutation in the second LIM domain of *FHL1* [21]. Because FHL2 contains the LIM domain, similar to FHL1 and MLP/CSRP3, it is reasonable to speculate that FHL2 is involved in autophagy to regulate the development of skeletal muscle. The present study examined this hypothesis.

## Materials and Methods

### Cell cultures

The mouse C2C12 myoblast cell line (Fuheng Cell Center, Shanghai, China) was maintained in growth medium composed of Dulbecco’s Modified Eagle Medium (DMEM), 10% fetal bovine serum (FBS), and 1% Antibiotic-Antimycotic (ABAM) at 37°C under 5% CO_2_. Differentiation into myotubes was activated by replacing the growth medium with differentiation medium composed of DMEM, 2% horse serum, and 1% ABAM.

### FHL2 silencing and overexpression

C2C12 cells were cultivated in 6-well plates and transfected with siRNAs (sense: 5’-GCAAGGACUUGUCCUACAATT-3’, antisense: 5’-UUGUAGGACAAGUCCUUGCTT-3’; Sangon Biotech, Shanghai, China) when grown to a density of approximate 70% in plates. In contrast, control cells were transfected with negative siRNA with same other condition. The transfection reagent was Lipofectamine 3000 (Invitrogen, Carlsbad, CA, USA). The knockdown efficiency was assessed by quantitative RT-PCR of FHL2 mRNA and western blot assay of FHL2 protein.

C2C12 cells were transfected with a plasmid pcDNA3.1 which was produced by cloning FHL2 cDNA into the pcDNA3.1 expression vector (Sangon Biotech, Shanghai, China). The transfection reagent Lipofectamine 3000 (Invitrogen) was mixed with optim-mem. Plasmid with optim-mem was mixed with Lip3000. Next, they were mixed all together at room temperature for 10 min. Finally, the mixture was added into C2C12 6-well plates at 37°C under 5% CO_2_ for 48h.

### RNA extraction, cDNA synthesis and RT-PCR

Total RNA was isolated using TRIzol (TAKARA, Dalian, China) reagent according to the manufacturers’ instruction. RNA was reverse transcribed by TAKARA PrimeScript^TM^ RT reagent kit (TAKARA) according to the manufacturers’ instruction. Quantitative RT-PCR assay was performed essentially as previously described [22].Primer are used in Table 1.

**Table 1.**
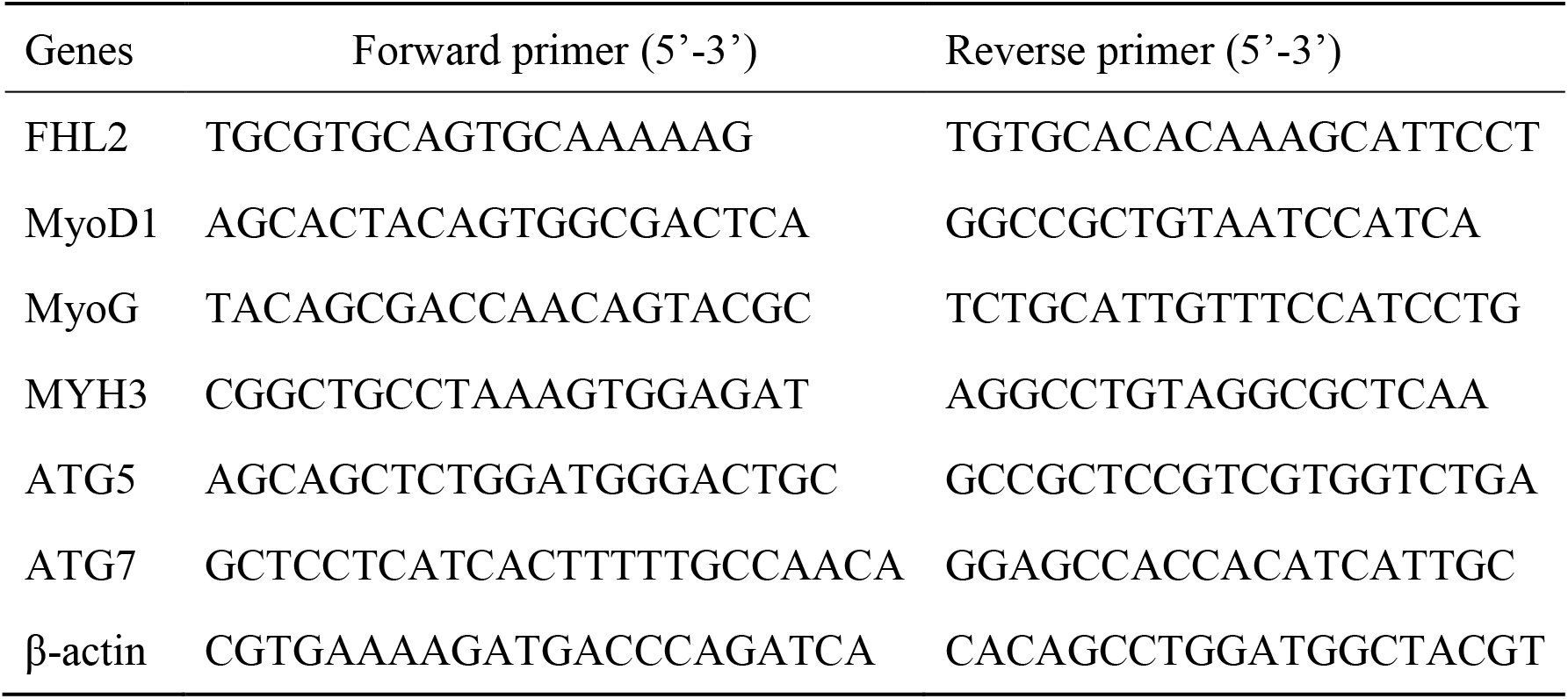
Primers used for quantitative real-time PCR

### Western blot assay

The cells were collected from the cultures, placed in the RIPA lysis buffer on ice (BestBio, Shanghai, China). The whole proteins were subjected to 10% sodium dodecyl sulfate polyacrylamide gel electrophoresis (SDS-PAGE) and then transferred to polyvinylidene fluoride membranes (PVDF; Millipore Corporation, Billerica, MA, USA). The PVDF membrane was incubated with 5% defatted milk powder at room temperature for 1 h, then incubation with the following specific primary antibodies at 4°C overnight: anti-FHL2 (Abcam, Cambridge, MA, USA), anti-MYOD1 (Abcam), anti-MyoG (Abcam), anti-MYH3 (Abcam), anti-ATG5 (Cell Signaling, Danvers, MA, USA), anti-ATG7 (Cell Signaling), and anti-β-Actin (Abcam). The secondary antibodies HRP-labeled mouse and rabbit IgG (Cell Signaling) were added at room temperature for 1h. Following each step, the membranes were washed five times with PBS-T for 3 min. The proteins were visualized by enhanced chemiluminescence (Amersham Pharmacia Biotech, Piscataway, NJ, USA) with a Kodak imager (Eastman Kodak, Rochester, NY, USA). Quantification of protein blots was performed using the Quantity One 1-D software (version 4.4.0) (Bio-Rad, Hercules, CA, USA) on images acquired from an EU-88 image scanner (GE Healthcare, King of Prussia, PA, USA).

### Microscopy

Cellular morphology was evaluated in proliferating myoblasts and differentiated myotubes by phase-contrast microscopy without preliminary fixation. Pictures were produced using the Olympus IX73 inverted microscope (OLYMPUS, Tokyo, Japan) and the Hamamatsu C11440 digital camera (HAMAMATSU, Shizuoka, Japan).

### Transmission electron microscopy

C2C12 myoblasts or myotubes were detached from the plates using a manual scraper, washed with PBS for a while. Then the cells were suspended and fixed overnight at 4 °C in 2% glutaraldehyde with 1% tannic acid in 0.1M sodium cacodylate, pH 7.3. The cells were rinsed three times in the sodium cacodylate buffer and incubated in 2% osmium tetroxide in the same buffer for 2 h at room temperature. Afterwards, the cells were rinsed three times in distilled water and exposed to 1% uranyl acetate in water for 15 min at room temperature and twice in distilled water. Next, the cells were spun down into 3% agarose at 45 °C and cooled to form blocks. The agarose blocks were dehydrated in graded steps of acetone and embedded in Spurr’s low-viscosity media. Following polymerization overnight at 65 °C, 80-nm sections were cut on a Reichert-Jung Ultracut E ultramicrotome and picked up on copper grids. The grids were post-stained in uranyl acetate and bismuth subnitrate. The sections were observed on a Philips CM-10 TEM (HT7700) and micrographs were recorded on a Kodak 4489 sheet film (Eastman Kodak).

### Immunofluorescence assay

Dry the cell climbing slides slightly and mark the objective area with liquid blocker pen for 20 min at room temperature, where add 50-100 μl of permeabilize working solution. Then eliminate obvious liquid, mark the objective tissue with liquid blocker pen. Cover objective area with 3% BSA at room temperature for 30 min and incubate cells with primary antibody overnight at 4°C, placed in a wet box. Cell climbing slides were covered with secondary antibody labelled with HRP at room temperature for 50 min. Then DAPI solution was added for incubation at room temperature for 10 min, kept in dark place. Put the slides on a glass microscope slide and then mount with resin mounting medium. Finally, Microscopy detection and images collection were performed by Fluorescent Microscopy (OLYMPUS).

### Co-immunoprecipitation assay

Protein concentration was determined using the BCA Protein Quantitation Kit (BestBio). 1mg of lysate was mixed with lysis buffer including phosphatase inhibitor to a volume of 1ml. Then Lysates were precleared with 5 μg of appropriate control IgG (Santa Cruz Biotechnology) and 20 μl of protein A/G plus-agarose (Santa Cruz Biotechnology) for 1 h rotation at 4 °C. Lysates were centrifuged (500 × g for 5 min at 4 °C) and 5 μg of FHL2 (Abcam) or LC3-II antibody (Abcam) or corresponding IgG was added to the precleared lysates and kept on ice for ~ 4 h. After incubation, 30 μl of protein A/G plus-agarose was added to each tube and kept on a rotator overnight at 4 °C. Lysates were then centrifuged (500 × g for 5 min at 4 °C). The pellet fractions were washed four times with PBS-PI and then resuspended in 20 μl of loading buffer. Samples were electrophoresed on a 12% SDS-PAGE gel and immunoblotted with the appropriate antibody.

### Flow cytometry assay

Apoptosis was induced by treating cells with 2uM Staurosporine (STS) (Selleckchem, Houston, TX, USA) in 12-well plates. Then cells were collected and digested by pancreatic enzyme to be cell suspension. Then the cells were washed twice with cold PBS and then resuspended cells in 1×binding buffer (BD Pharmingen, Santiago, CA, USA) at a concentration of 1×106 cells/mL. One hundred microliters of the solution were transferred to a 5-mL culture tube, and then 5 μL of Annexin V-FITC (BD Pharmingen) and 5 μL of PI (BD Pharmingen) were added. The cells were gently vortexed and incubated for 15 min at room temperature (25°C) in the dark. Four hundred microliters of 1×binding buffer was added to each tube and analyzed by flow cytometry (BD FACSCalibur, BD Pharmingen) within 1 h.

### Statistical analysis

All statistical analyses were performed using SPSS 17.0 (SPSS Inc., Chicago, IL, USA). Data are presented as least squares means ± standard error of the mean (SEM), and values were considered statistically different at *P*<0.05.

## Results

### The role of FHL2 in skeletal muscle differentiation

To explore the potential role of FHL2 in skeletal muscle differentiation, we performed a knockdown assay in C2C12 myoblasts derived from mouse satellite cells. C2C12 cells transfected with FHL2 siRNA or scrambled siRNA were induced to differentiate, and *FHL2* mRNA expression was shown to be reduced significantly after knockdown in both myoblasts and myotubes compared with controls (Fig. 1A). Western blot analysis revealed a decrease in FHL2 protein in FHL2-silenced cells compared with control cells (Fig. 1B). Next, morphological differences between negative control and FHL2 siRNA-transfected groups were compared during C2C12 differentiation into myotubes. The FHL2-silenced group showed reduced myotube formation (Fig. 1C), and the expression of myogenic marker genes *MyoD1*, *MYH3*, and *MyoG* was significantly reduced in FHL2-silenced cells compared with controls (Fig. 1D–F). Moreover, western blotting revealed that MYH3 and MyoG protein levels were reduced after FHL2 silencing (Fig. 1G). These results suggest that FHL2 siRNA was effective and that FHL2 plays an important role in muscle differentiation by regulating myogenesis-related genes. The overexpression of FHL2 in C2C12 myoblasts and myotubes (Fig. 2A and B) had no significant effect on *MyoG*, *MyoD1*, or *MYH3* expression levels (Fig. 2C–E).

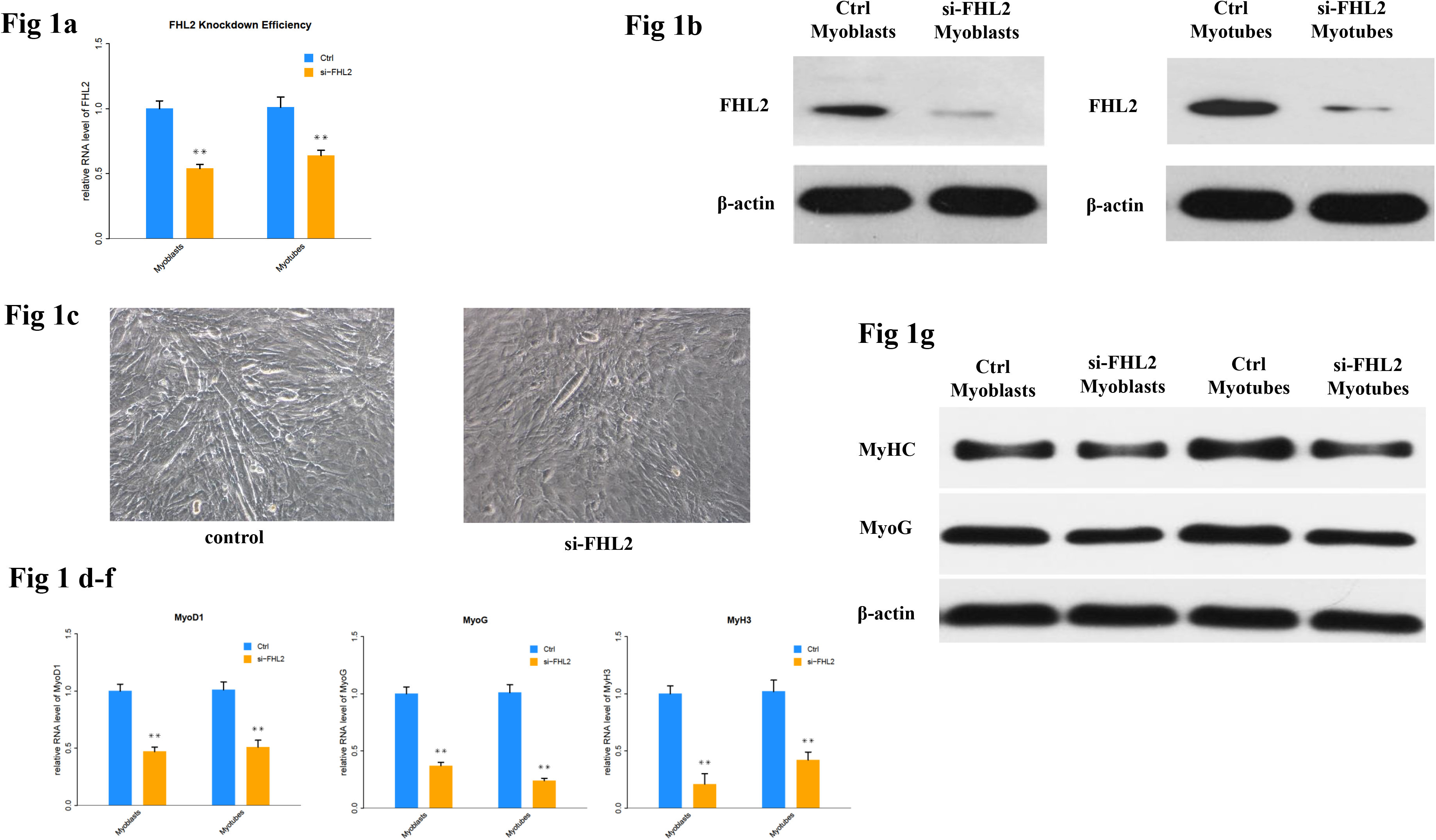

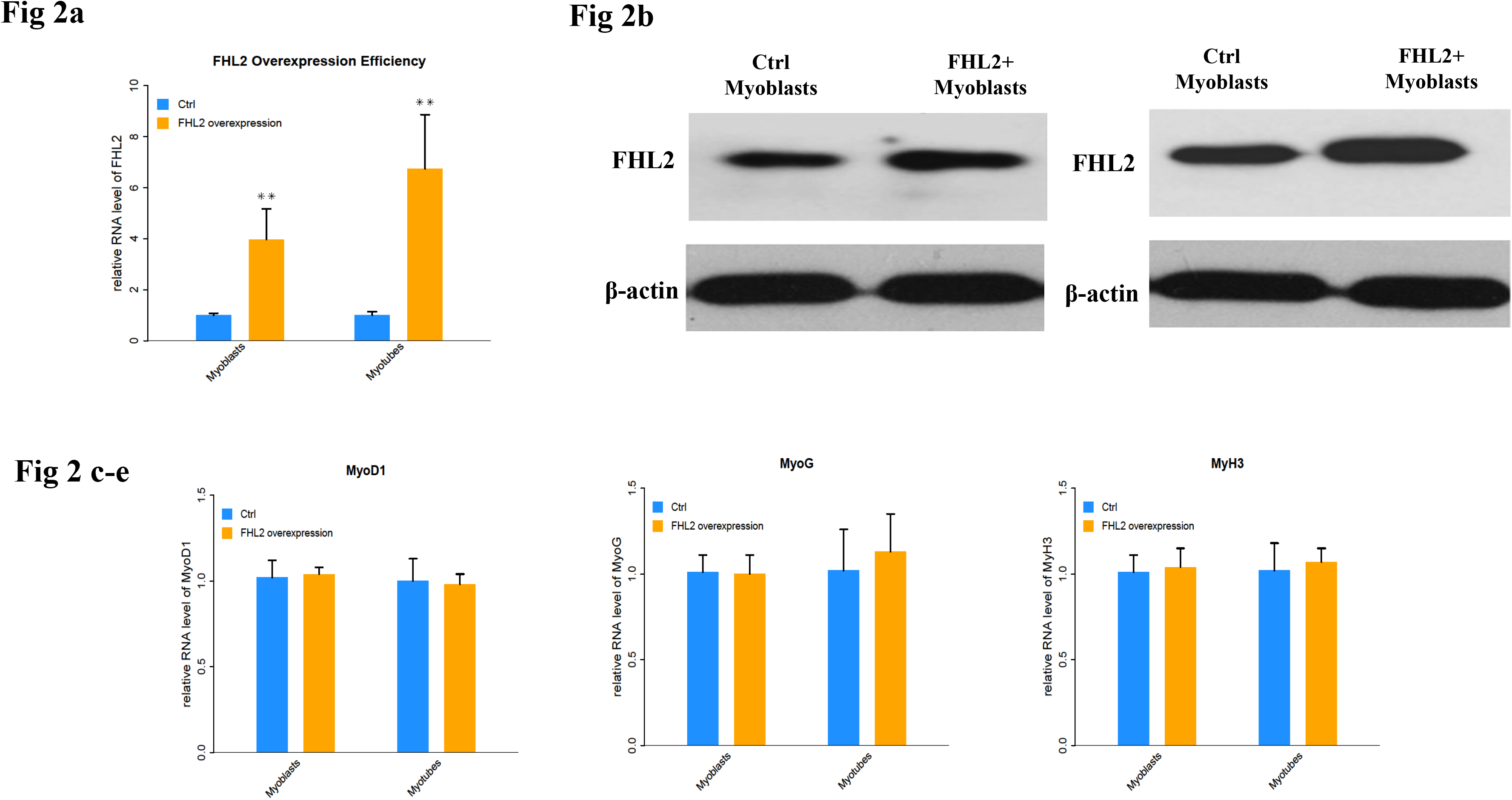

### FHL2 regulated autophagy in skeletal muscle cells

To determine whether FHL2 silencing in skeletal muscle influenced the induction of autophagy, the expression of autophagy genes *ATG5* and *ATG7* was measured and shown to be significantly reduced in myoblasts and myotubes from FHL2-silenced cells compared with control cells (Fig. 3A). Next, LC3 protein level changes were examined to monitor autophagy induction, and the ratio of LC3-Ⅱ to LC3-I protein was found to be reduced in FHL2-silenced myoblasts and myotubes compared with controls (Fig. 3B). Interestingly, the starvation of FHL2-silenced myoblasts and myotubes, which should have been able to activate autophagy, did not induce the accumulation of LC3-Ⅱ (Fig. 3B). Moreover, FHL2 overexpression did not significantly influence the expression of *ATG5* or *ATG7* (Fig. 3C).

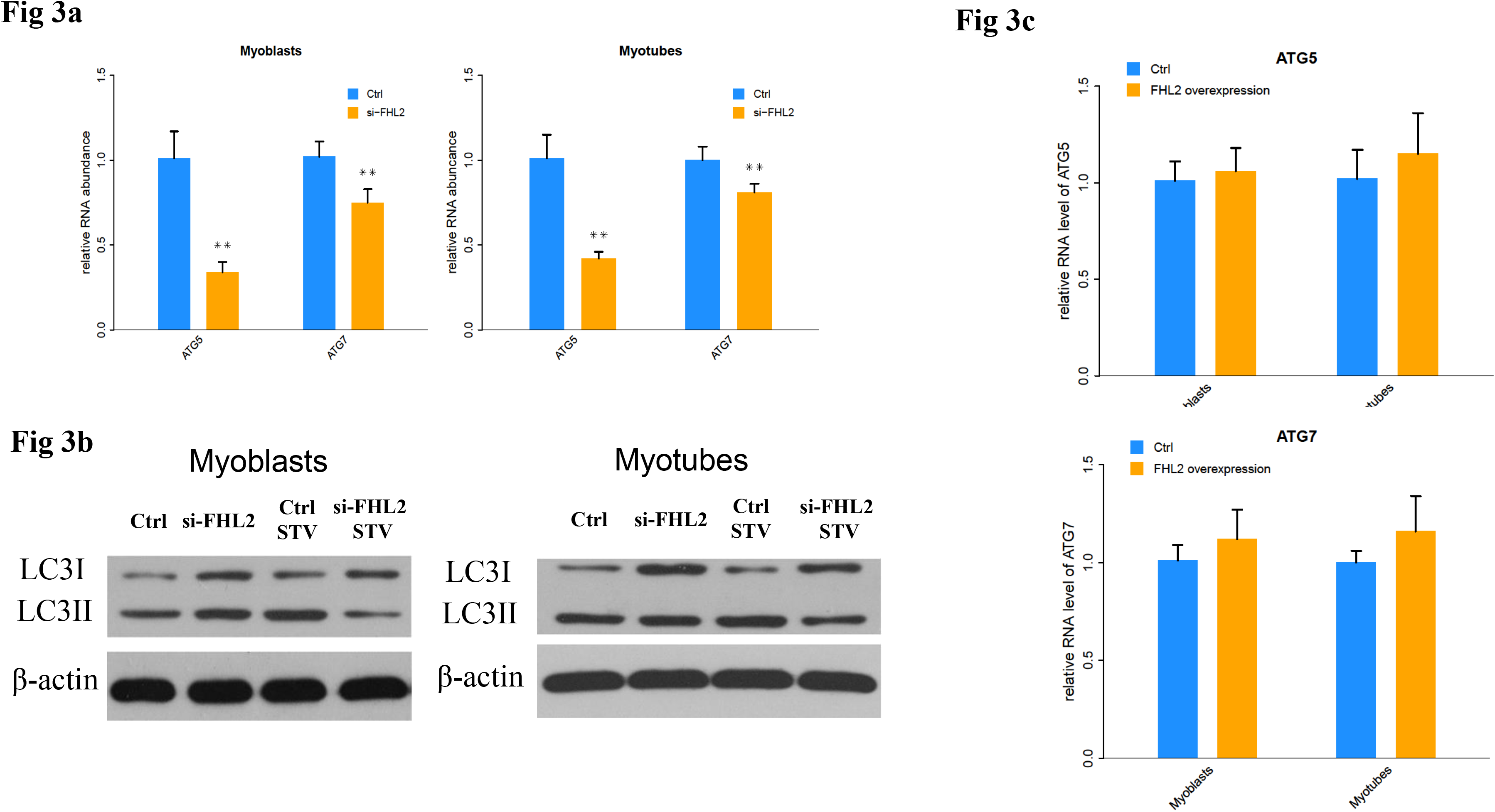

To further confirm our findings, we used TEM to observe the ultrastructure of myoblasts. As shown in Fig. 4, both negative control and FHL2-overexpressing myoblasts were observed to contain normal autophagosomes whereas large empty autophagosomes were present in FHL2-silenced myoblasts, which was indicative of impaired autophagy.

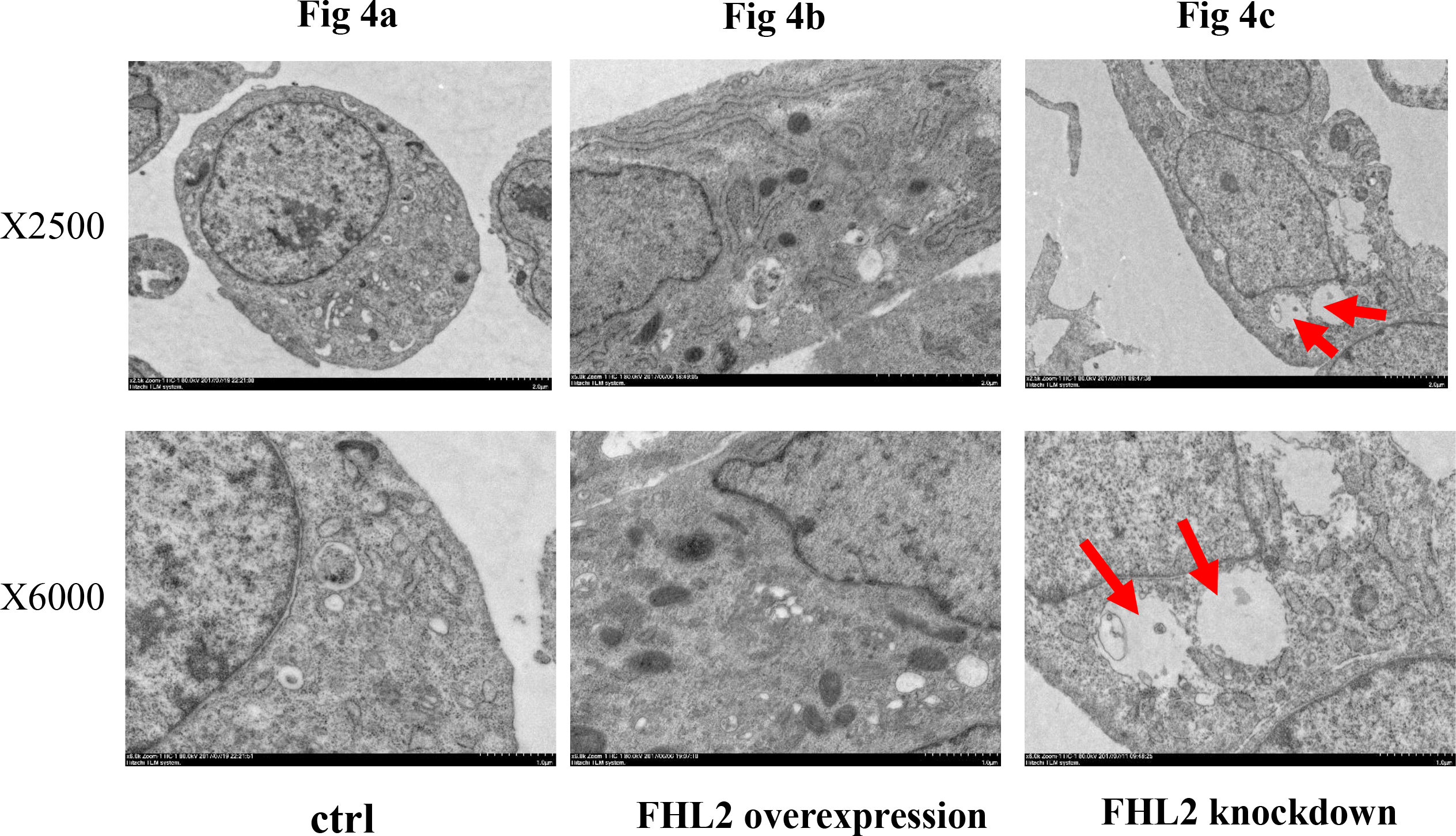

### FHL2–LC3-Ⅱ co-localization

Immunofluorescence and co-immunoprecipitation assays were next performed to determine whether FHL2 co-localized with the autophagosomal marker LC3 during myoblast differentiation. Immunofluorescence analysis showed similar FHL2 and LC3 protein distribution in myoblasts (Fig. 5A), while FHL2 silencing decreased both FHL2 and LC3Ⅱ expression which was indicative of their co-expression (Fig. 5A). The co-immunoprecipitation assay revealed co-localization of FHL2 and LC3 proteins in C2C12 cells during myoblast differentiation, and reduced co-immunoprecipitation in FHL2-silenced myoblasts compared with control cells (Fig. 5B). These results support a role for FHL2 in the formation of autophagosomes by binding LC3 in an intracellular complex.

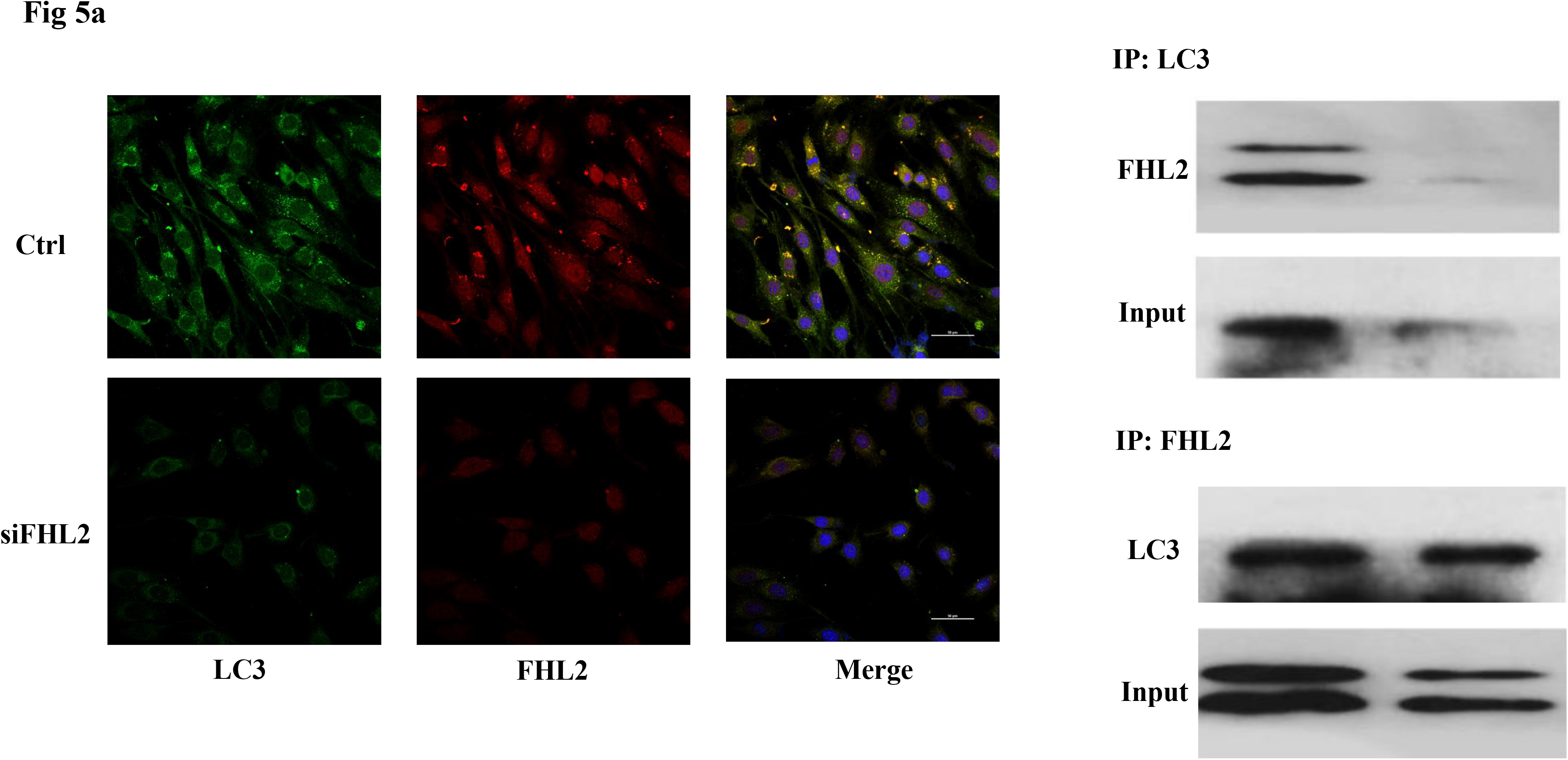

### FHL2 silencing promoted apoptosis in both myoblasts and myotubes

Flow cytometry was next used to measure cell death after treatment of C2C12 cells with 2 μM STS for 12 h. Cell death was observed in 2.74% and 3.51% of wild-type (WT) myoblasts and myotubes, respectively, compared with 6.30% and 10.77%, respectively, in si-FHL2 groups (Fig. 6B and D). This indicated that FHL2 silencing enhanced apoptosis in both myoblasts and myotubes. Moreover, a decreased resistance to STS was detected in FHL2-silenced myoblasts and myotubes (Fig. 6A–D). Western blotting of PARP and caspase-3 proteins under different conditions revealed the increased accumulation of PARP in myoblasts following FHL2 silencing (Fig. 6E). Caspase-3 protein expression was not detected in WT cells, but was observed in si-FHL2 myotubes, suggesting an important role for FHL2 in protecting myotubes from apoptosis (Fig. 6E). We also found that myotubes were more resistant to STS-induced apoptosis than myoblasts (Fig. 6E).

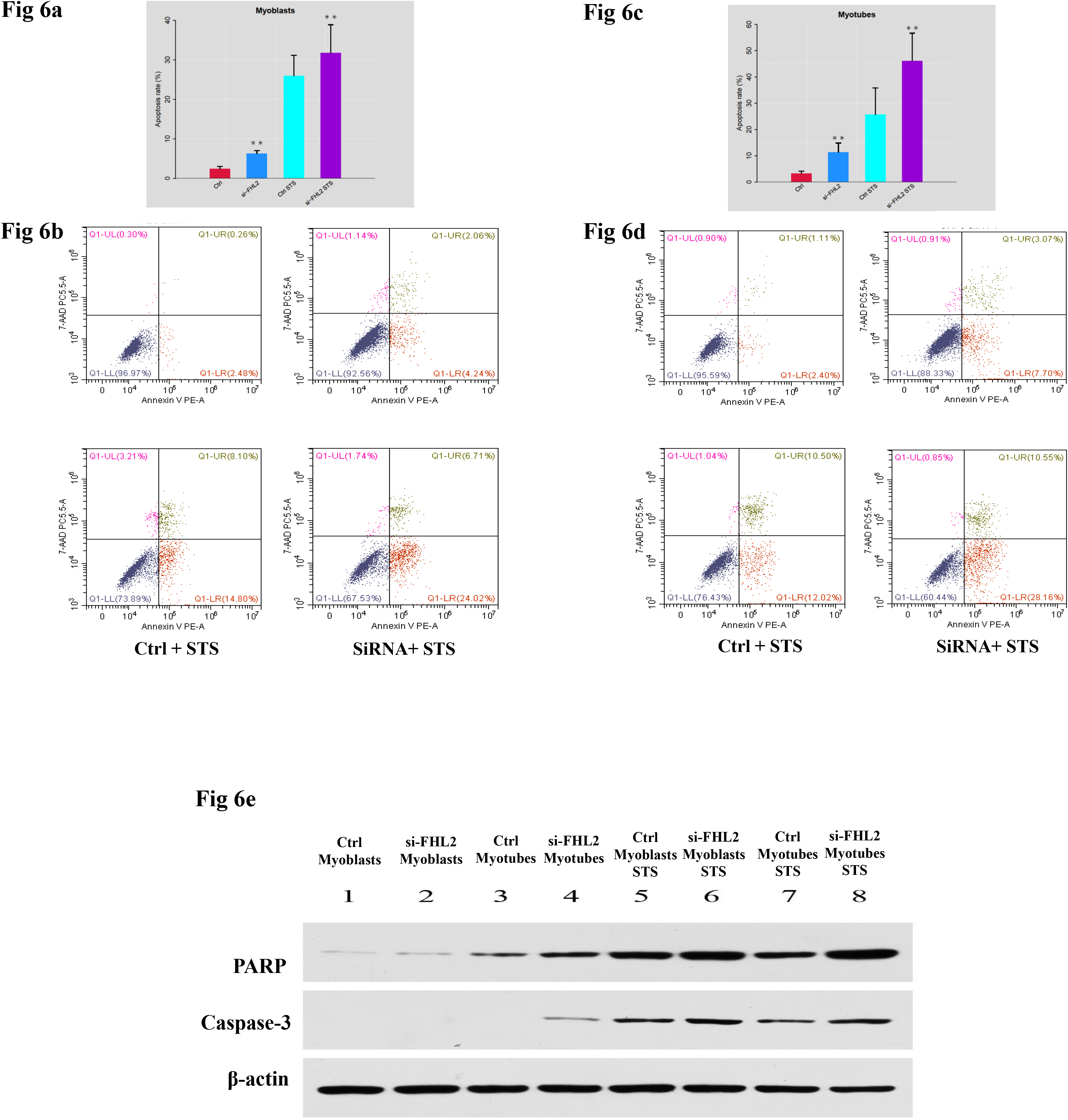

## Discussion

Although previous studies suggested that FHL2 is involved with autophagy and may be associated with LC3 protein [19, 23], its autophagy-related role in muscle differentiation had not been fully determined. In this study, we explored the mechanism of FHL2 in muscle differentiation through autophagy induction.

We initially confirmed that FHL2 played a role in muscle development by measuring the gene and protein expression of muscle-related MyoD1, MyH3, and MyoG after FHL2 silencing and overexpression. FHL2 expression was significantly lower after siRNA transfection, indicating the efficiency of siRNA. Expression of the muscle-related genes was decreased in FHL2-silenced myoblasts or myotubes, and their protein expression was slightly reduced. However, no reduction in their expression was detected in FHL2-overexpressing cells. Those results confirm that FHL2 play a part in muscle development.

Previous studies detected LC3 protein accumulation during muscle cell development by western blotting [24, 25]. We investigated the relationship between FHL2 and autophagy using two methods: first, measuring the expression of genes essential for autophagy such as *ATG5* and *ATG7* [26] following FHL2 knockdown, and second, measuring the correlation between FHL2 and LC3Ⅱ. Knockdown, but not the overexpression, of FHL2 significantly decreased the expression of *ATG5* and *ATG7* in both myoblasts and myotubes. Additionally, the autophagy level as indicated by the ratio of LC3-Ⅱ to LC3-I protein [27] was reduced in FHL2-silenced myoblasts and myotubes. These results indicate a role for FHL2 in autophagy induction.

It is reasonable to assume that the function of FHL2 in muscle development is associated with autophagy because similar effects on muscle-related genes and autophagy markers were detected following FHL2 silencing or overexpression. Moreover, the starvation that activated autophagy in control cells did not induce LC3-Ⅱ accumulation in either FHL2-silenced myoblasts or myotubes, indicative of a function for FHL2 in autophagy induction. Further TEM analysis showed that FHL2 knockdown caused organelle abnormalities and impaired autophagy in C2C12 cells, but not in control or FHL2-overexpressing C2C12 cells. The persistent activation of catabolic pathways in muscle causes atrophy and weakness, and autophagy is required to maintain muscle mass [17, 28]. Therefore, our results suggest that FHL2 has a role in myotube formation by activating cell autophagy to maintain cellular homeostasis.

MLP/CSRP3 containing a LIM domain has been reported to participate in muscle differentiation and autophagosome formation by interacting with LC3-Ⅱ [20]. FHL2 also contains four and a half LIM domains that act as a protein–protein binding interface involved in muscle differentiation [29]. Considering the potential of FHL2 to bind LC3-Ⅱ, its correlation with LC3-Ⅱ at the protein level in C2C12 cells, and its likely role in the assembly of extracellular membranes [30], we hypothesized that FHL2 participates in autophagosome formation by interacting with LC3-Ⅱ. We therefore next explored their localization in C2C12 cells. The protein distribution of FHL2 in myoblasts was shown to resemble that of LC3-Ⅱ, and immunoprecipitation revealed the co-localization of LC3-Ⅱ and FHL2. More importantly, FHL2 silencing simultaneously reduced both FHL2 and LC3 protein expression. These findings provided evidence of a role for FHL2 in muscle development by interacting with LC3-Ⅱ and participating in the formation of autophagosomes.

Insufficient autophagy contributes to the accumulation of waste material inside the cell and the induction of apoptosis. However, autophagy can also protect cells from apoptosis [31]. Study has revealed autophagy protects myoblasts against apoptosis [32]. In a previous study, the specific autophagy inhibitor 3MA [33] was used to treat C2C12 cells, resulting in increased *CASP3* activity which is a marker of increased apoptosis [18]. If FHL2 functions in the induction of autophagy, FHL2-silenced C2C12 cells would be expected to show more apoptosis. Indeed, we observed increased STS-induced death of both FHL2-silenced myoblasts and myotubes compared with controls. These results suggested that FHL2 knockdown reduces autophagy, indicating that it regulates muscle development by controlling autophagy and thereby FHL2 may regulate muscle development through autophagy against apoptosis. However, exactly how autophagy prevents apoptosis requires additional study. Furthermore, to validate the results of flow cytometry, the protein levels of cleaved caspase-3 and PARP-1, markers of apoptosis [34, 35], were measured in WT and STS-treated groups by western blotting. Increased cleavage of PARP-1 and caspase-3 was observed in FHL2-silenced cells relative to controls, and increased PARP-1 cleavage was also detected during the differentiation of FHL2-silenced cells. These results reflect increased apoptosis in the absence of FHL2, likely caused by repressed autophagy.

In conclusion, our findings suggest that FHL2 interacts with LC3-Ⅱ protein to regulate muscle development through autophagy. As a result, the deletion of FHL2 inhibited muscle development and damaged autophagosomes. However, further studies of mechanism of autophagy in muscle cells are needed to understand in detail how FHL2 regulates autophagy and its contribution to muscle development or myopathies.

## Acknowledgments

This work was financially supported by the China Agriculture Research System (CARS-40), and the Thirteenth Five Year Plan for Breeding Program in Sichuan (2016NYZ0050). We thank Sarah Williams, PhD, from Liwen Bianji, Edanz Group China (www.liwenbianji.cn), for editing the English text of a draft of this manuscript.

